# The hidden prevalence and unique protein folds of huge phages

**DOI:** 10.1101/2025.02.27.640493

**Authors:** LinXing Chen

**Affiliations:** State Key Laboratory of Advanced Environmental Technology, the Department of Environmental Science and Engineering, University of Science and Technology of China, Hefei, China, 230026

**Keywords:** viruses, huge phages, jumbo phages, metagenomics, microbial ecology, protein structures

## Abstract

Huge phages are remarkable for their expansive genomes (≥200 kbp, and up to 841 kbp to date), global distribution, and intricate functional repertoires. However, their study has been hindered by challenges in genome identification and the lack of a comprehensive reference database. Using vRep, we established the Huge Phage Genome Collection (HPGC), a curated resource of 7,295 non-redundant huge phage genomes spanning 4,613 species-level groups. Within HPGC, we identified an overlooked and prevalent clade, HP8026, which constitutes ~1.2% of human gut metagenomic reads, suggesting its potential in fecal contamination monitoring and a promissing modulator of gut microbiome dynamics. Complementing HPGC, we developed the Huge Phage Protein Structure Database (HPDB), which catalogs 15,594 structure clusters derived from 589,960 predicted proteomes. Notably, 40% of these clusters presented in huge phages only, exhibit specific protein folds. Additionally, 3% of the HPDB clusters were absent in other viral forms, including the plasmid-originated ArdC that could protect huge phages from the antiviral mechanisms of their hosts. Collectively, HPGC and HPDB provide a pioneering framework for exploring the evolution, functional innovation, and biotechnological potential of huge phages, while serving as indispensable references for viral ecology and structural biology.

## Introduction

Viruses are the most abundant biological entities on Earth ^1^, and those specifically infect bacteria are known as bacteriophages (or phages). Phages are widely distributed in various natural and host-associated habitats ^2^. They could dramatically influence microbial communities by infecting and killing bacterial hosts, altering their metabolic potentials via auxiliary metabolic genes (AMGs), and driving their evolution via horizontal gene transfer. The double-strand DNA phages usually have a genome size of approximately 50 kbp ^3^, while some of them, named jumbo phages or huge phages, are tailed phages with genomes exceeding 200 kbp in length ^3–5^.

Huge phages are diverse and widespread, their huge genome sizes (up to 841 kbp ^6^) and intricate functional repertoires have thus provided a unique opportunity to study more complex biological mechanisms in viruses. Huge phages were initially studied via isolation, yet have been challenging due to their large virions, which block diffusion in semisolid media, preventing the formation of visible plaques ^7^. Thus, genome-resolved metagenomics has been the primary approach ^3,8–17^. Huge phages encode tens of tRNAs and have genes for functions related to transcription and translation ^3^, making them distinguished from small phages. Some huge phages are equipped with CRISPR-Cas systems to confer their hosts with extra antiviral weapons ^15^. Besides, some freshwater huge phages infecting aerobic methane oxidizers could be directly involved in methane oxidation via their pmoC, an enzymatically critical subunit of the particulate methane monooxygenase ^9^. More interestingly, some huge phages could amplify their DNA in a nucleus-like structure without being targeted by the CRISPR-Cas systems of their hosts ^18–21^. Despite these advances, the misidentification of huge phage genomes in metagenomic analyses, the lack of a reliable reference genome database, and the difficulties in annotating most protein-coding genes hinder comprehensive insights into their diversity, distribution, and ecological significance.

In this study, we developed the vRep analysis pipeline to identify viral genomes with high confidence from metagenomic assemblies. We collected 10,690 potential huge phage sequences from public databases and publications (before 30 September 2024), and constructed the curated Huge Phage Genome Collection (HPGC) with vRep. HPGC contained 7,295 non-redundant genomes comprising 4,613 species-level groups, including the HP8026 clade that predominates in many human gut metagenomes. Protein structure analysis of HPGC genomes with the Huge Phage protein structure DataBase (HPDB) suggested the prevalence of protein structures unique to huge phages while not in other viral equivalents like small phages and eukaryotic viruses. Together, huge phages could be key in both host-associated and natural habitats, and the construction of HPGC and HPDB will accelerate the study of huge phages.

## Results

### The identification of viral sequences with vRep

The initial and crucial step in metagenomics-based viral research is the identification of viral genomic sequences. Various tools are available for this purpose, including Virfinder ^22^, Virsorter2 ^23^, VIBRANT ^24^, CheckV ^25^, and geNomad ^26^. Often, researchers employ multiple tools concurrently in their studies. However, it is generally preferable to verify whether a sequence is viral using a single tool rather than relying on the consensus of multiple tools, as this approach tends to yield a larger number of viral sequences. Despite the fact these tools have largely enlarged our knowledge of viruses, it is important to recognize that they are not perfect, and false positives are known to occur in published databases.

To address the challenges of identifying viral sequences from metagenomic assemblies, we developed a pipeline named vRep, which integrates geNomad, VirSorter2, and CheckV (Figure 1). The core concept of vRep is to leverage multiple tools for enhanced accuracy in viral identification. Initially, only the sequences identified as virus-containing (provirus or full-length virus) by both geNomad and VirSorter2 are processed through CheckV to remove host fragments. If a sequence was flagged as “provirus” by both geNomad and VirSorter2 but “full-length viruses” by CheckV, it will be excluded from subsequent analyses. Next, the purely viral sequences without host fractions will undergo a second evaluation with VirSorter2 without predicting proviruses. Next, the information from CheckV and the second VirSorter2 run on the purely viral sequences is used to refine the selection. A sequence that meets any of the following criteria will be retained: (1) it contains at least one viral gene; (2) it lacks identified viral genes but also (a) has no host genes, or (b) possesses a viral score of ≥ 0.95, or (c) has ≥ 2 viral hallmark genes. Subsequently, the retained sequences are assessed for potential large genome duplication issues during assembly (see below for details). This issue could be detected by CheckV ^25^ via Kmer frequency calculation and evaluation. Ultimately, vRep generates a comprehensive summary file detailing all identified viral sequences, including their length, key data from geNomad, CheckV, and VirSorter2, as well as any detected duplication issues. Detailed parameters for each stage are specified in the Methods section.

**Figure 1.**
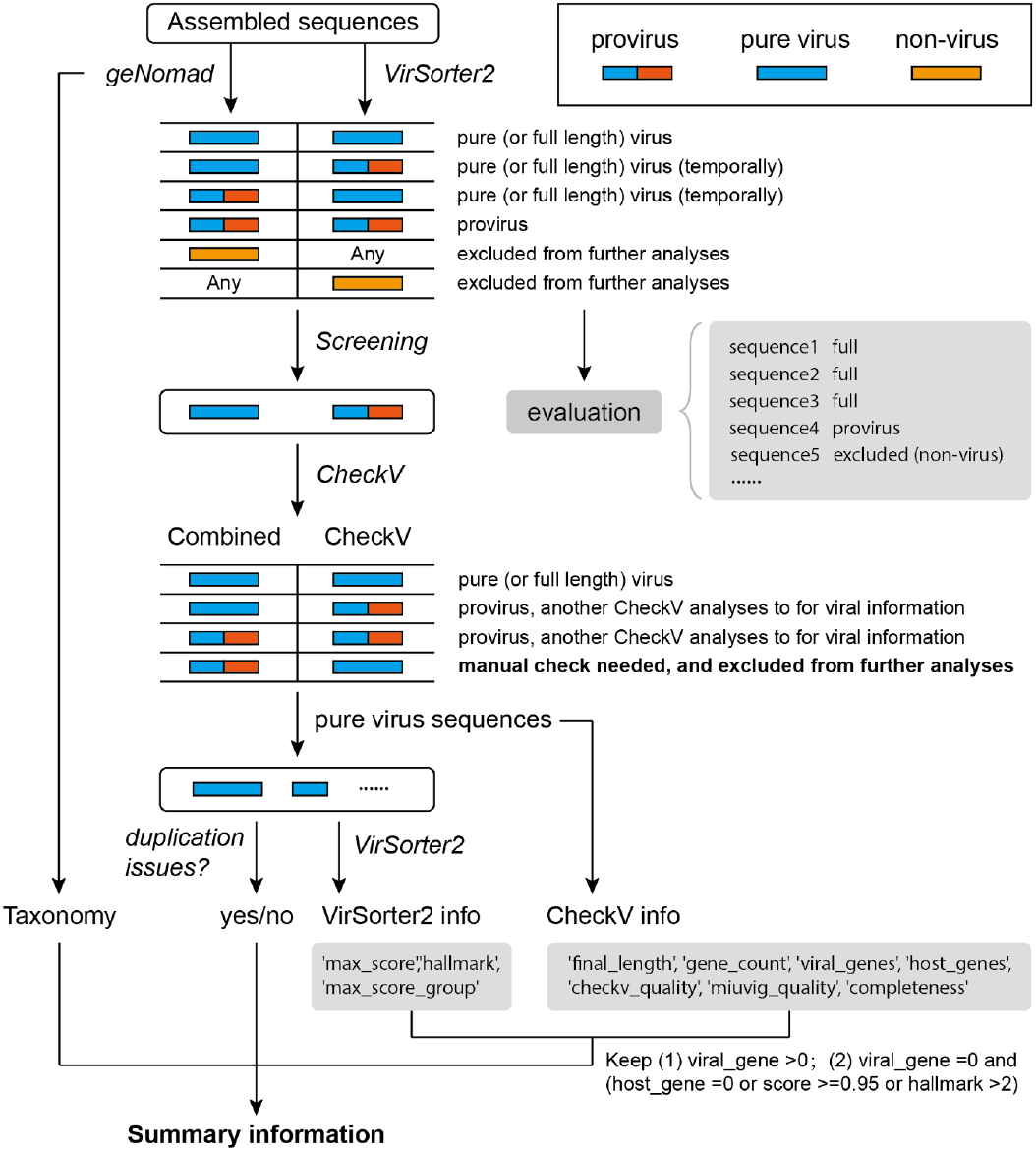
Viral sequence identification from metagenomes using vRep. The analyses start from assembled sequences, first predicted virus-containing sequences using geNomad and VirSorter2. If one or both tools predict a given sequence as non-viral, it will be excluded from further analyses. The identified viral sequences are then analyzed using CheckV to remove the host fractions. If a sequence was identified as a provirus by geNomad and VirSorter2 but a full-length virus by CheckV, a manual check will be asked. The pure viral sequences are subsequently evaluated by VirSorter2 again. The information from CheckV and the second VirSorter2 run is used to screen for viral sequences with high confidence. Also, a BLASTn search was performed to exclude duplication issues in metagenomic assembly.

### The misidentification of huge phage genomes is not rare

Our routine analyses revealed that some genomic sequences could be misidentified as huge phage genomes for various reasons that vRep could identify (Figure 2a). One common scenario involves prophage and their host genomic fractions side by side (case I). Occasionally, multiple prophages within a small region of the host chromosome can confuse phage identification tools, though such cases are relatively rare. Another issue arises when sequences from non-viral genetic elements (case II), or NCLDVs (nucleocytoplasmic large DNA viruses or giant viruses) (case III), are mistakenly classified as huge phage genomes. Lastly, as previously reported ^8^, some assemblies may generate sequences containing multiple copies of a specific genomic region where the length of a single genome copy is less than 200 kbp (case IV).

**Figure 2.**
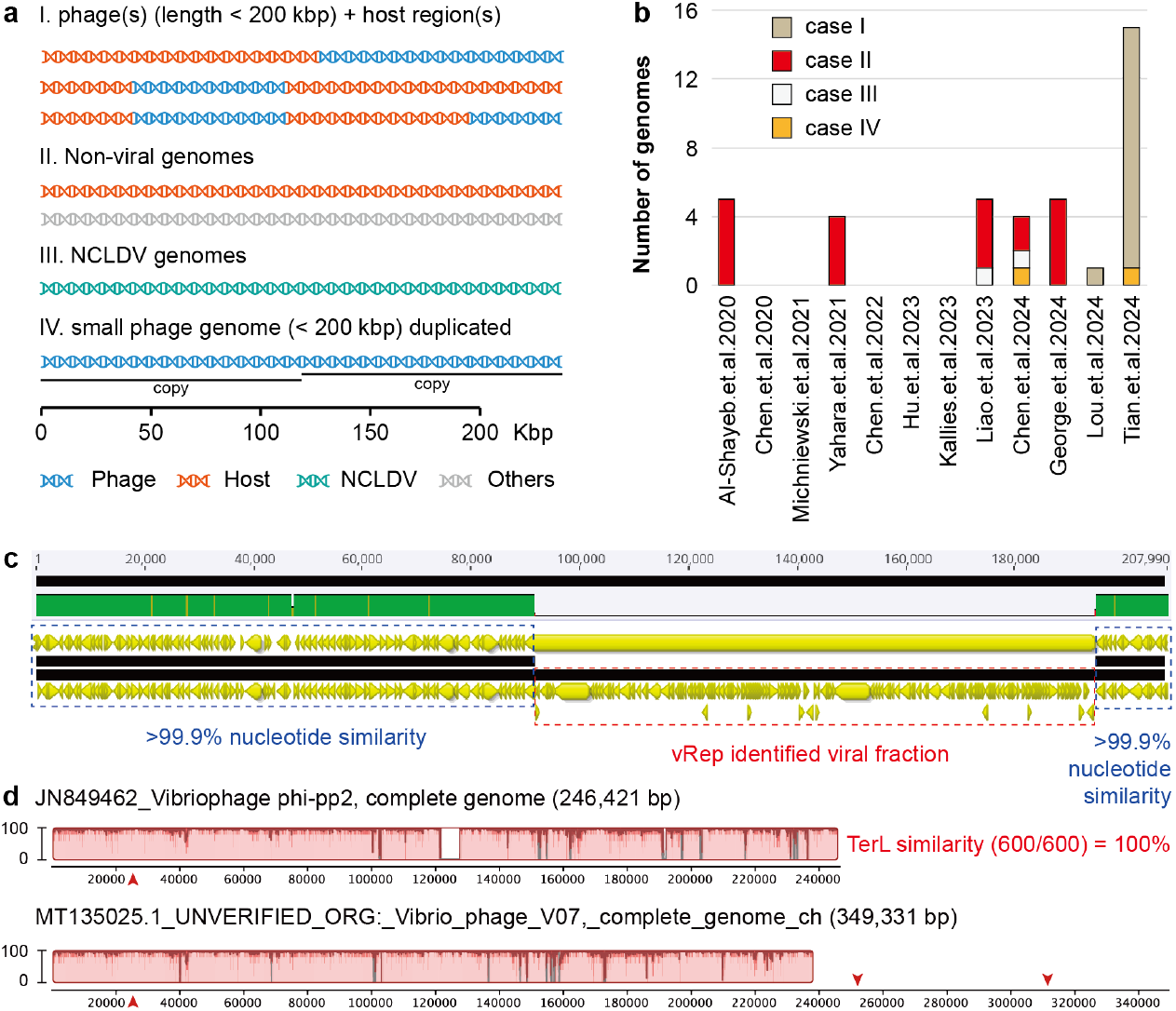
The misidentification of huge phage genomes in metagenomic analysis. (a) Illustration of examples in which different genomic resources may be identified as huge phage genomes. NCLDVs, nucleocytoplasmic large DNA viruses, or giant viruses. (b) The number of misidentifications of huge phage genomes in the publications focusing on phages. (c) An example of the case I issue of huge phage genome misidentification. The problematic huge phage genome contains a provirus (~103 kbp; red box) and its host genome (blue boxes). (d) An example of several concatenated phage genomes into a longer one. The problematic genome (bottom) and a complete reference genome (upper) are shown, with their pairwise genomic similarity indicated by the red profiles (range from 0 to 100). The red arrows indicate the locations of the large terminus encoding genes.

To assess whether these misidentification issues are common in studies of huge phages, we examined viral sequences ≥ 200 kbp from several publications focused on phages (Supplementary Table 1). From this analysis, we collected and evaluated 1,211 potential huge phage genomes from 12 publications. Among these, 39 genomes (~3.2%) were found to be misclassified, and 13 (~1.1%) could not be determined (Figure 2b). Of these misclassified genomes, 15 exhibited the case I issue (see Figure 2c and Supplementary Figure 1 for examples), which can lead to the identification of numerous false-positive auxiliary metabolic genes (AMGs) and result in inaccurate interpretations of the ecological roles of the corresponding phages. The most frequent issue, Case II, involved 20 genomes. This issue is prevalent not only in large phage studies but in broader viral research as well. As for case III, we found 2 NCLDV genomes were misidentified as huge phages. Although only 2 genomes from these phage-focused publications showed large fragment duplication (case IV), more instances of this issue were identified in other publications or databases that encompass all viruses rather than focusing solely on phages (Supplementary Figure 2).

The evolution of huge phages presents an intriguing and unresolved question: did huge phages evolve from smaller predecessors, or did they originally emerge with large genome sizes ^3,27^? We have identified an additional complication that vRep could not detect, and one might incorrectly conclude that huge phages evolved from smaller genomes without recognizing these misassemblies. This complication arises from the misassembly of two or more viral genomes into a single sequence, which may be detected, in some cases, by using single-copy viral marker genes such as the large terminase (terL) (Figure 2d).

### The huge phage genome collection, HPGC

Through vRep-based evaluation and filtering of the 10,690 potential sequences, we identified 9,372 huge phage genomes, ultimately establishing a non-redundant dataset of 7,295 high-quality genomes (Supplementary Tables 2). This curated Huge Phage Genome Collection (HPGC) represents the most comprehensive repository of huge phage genomes to date.

The non-redundant genomes originated from diverse ecological niches, with predominant contributions from animal-associated environments (31.4%, 2,288 genomes) and freshwater ecosystems (25.8%, 1,882 genomes). Additional sources included soil (12.8%, 936 genomes), marine systems (9.7%, 708 genomes), deep subsurface environments (4.7%, 345 genomes), and other habitats (Figure 3a). The genome size distribution analysis revealed a strong bias toward 200-400 kbp genomes (94.3%, 6,877 genomes), with only rare instances exceeding 600 kbp (Figure 3b). Notably, 69.9% of the genomes met MIUViG high-quality standards (Figure 3c).

**Figure 3.**
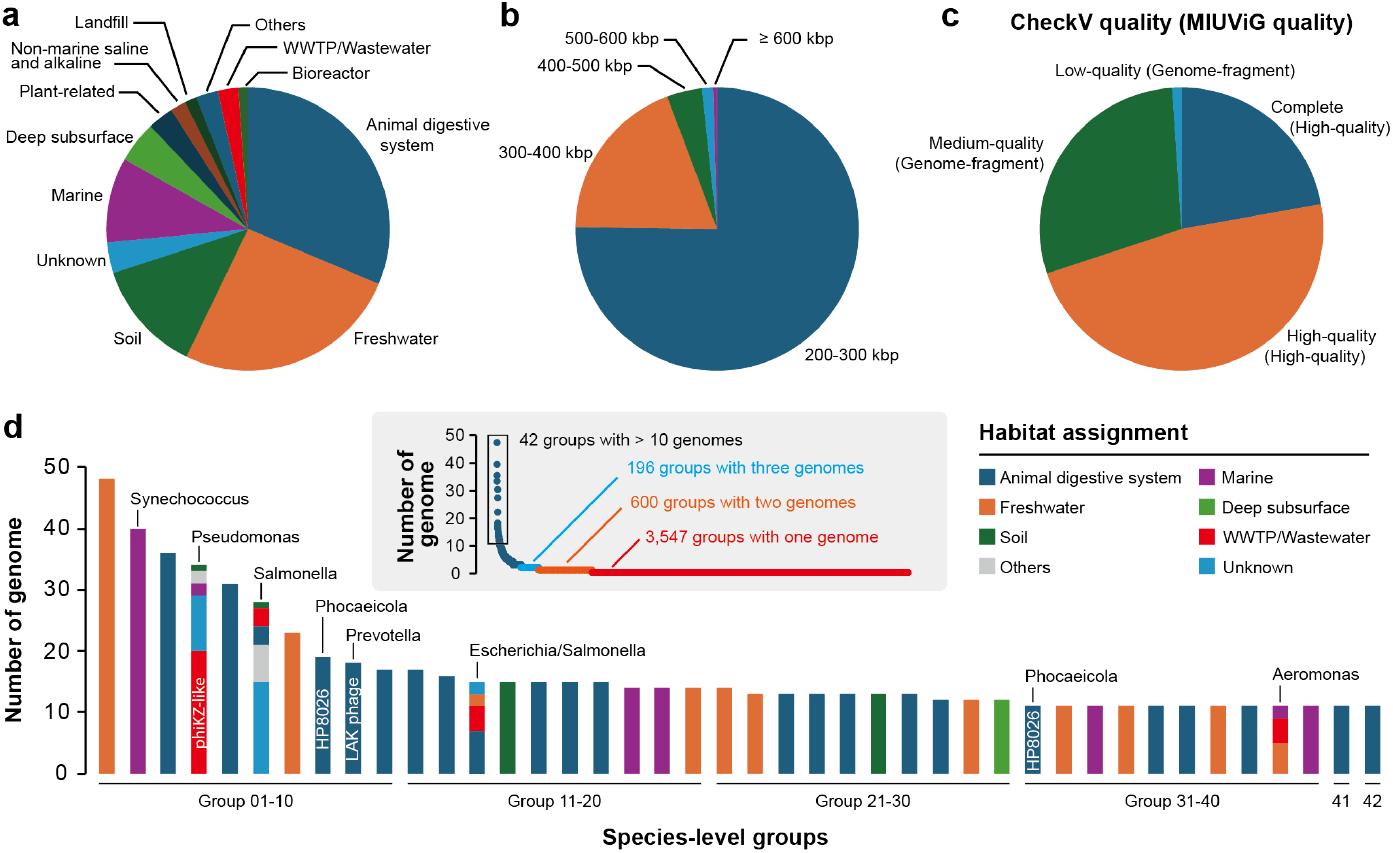
The general information of the 7,295 non-redundant huge phage genomes. (a) The habitat distribution. Only those habitats that account for 1% or more of the non-redundant genomes are shown, otherwise assigned to “Others”. Those without habitat information were assigned to “Unknown”. (b) The length distribution. (c) The genome quality evaluated by CheckV. (d) The species-level groups that contain > 10 genomes. The insertion shows the rank distribution of all species-level groups, with the number of groups containing one genome, two genomes, and three genomes shown. The bacterial hosts and the phage clade of the groups are shown when available.

Clustering analysis at 95% nucleotide identity delineated 4,613 species-level groups, highlighting the substantial genomic diversity of huge phages. Most groups (3,547) were singletons, and only 600 groups (13.0%) contained two genomes (Figure 3d). This diversity pattern persisted at lower identity thresholds, with 4,282 groups defined at 90% and 4,205 groups at 80% nucleotide identity. Notably, 1,173 (25.4%) and 2,177 (47.2%) of the group representatives were complete and high-quality, respectively, and only 54 representatives were evaluated as low-quality (Supplementary Table 2).

For the 42 species-level groups with > 10 genomes, habitat specificity was observed in 38 of them. Cross-environment distribution was limited to several groups: the extensively studied phiKZ-like phages targeting *Pseudomonas* spp., the LAK phages with *Prevotella* spp. as host, and those infecting *Synechococcus, Phocaeicola Salmonella, Escherichia*, and *Aeromonas* species.

### HP8026 is a clade widespread in human guts revealed by HPGC

One benefit HPGC offers is identifying the overlooked but prevalent huge phage clades. A cluster comprising 94 genomes was identified when clustering the non-redundant genomes at 80% sequence similarity (HP8026 for short hereafter) (Supplementary Table 3). They were assigned to 22 species-level groups, including groups 08 and 31 (Figure 3d). The other reasons for clustering them together include their high similarity of large terminases (> 99%; Supplementary Figure 3) and the sharing of a large number of protein clusters (Supplementary Figure 4).

Interestingly, all the HP8026 genomes were reconstructed from human gut samples collected in China, the USA, Spain, and other countries (Figure 4a). They have a length of 201-401 kbp and a GC content of 29.3-30.4% (Figures 4b and c). Eight genomes were predicted to be complete with an average genome size of 395 kbp (384-401 kbp). We manually curated all the 11 genomes shorter than 220 kbp to a final average length of 395 kbp (Figure 4b), suggesting that HP8026 genomes are around 400 kbp in size.

**Figure 4.**
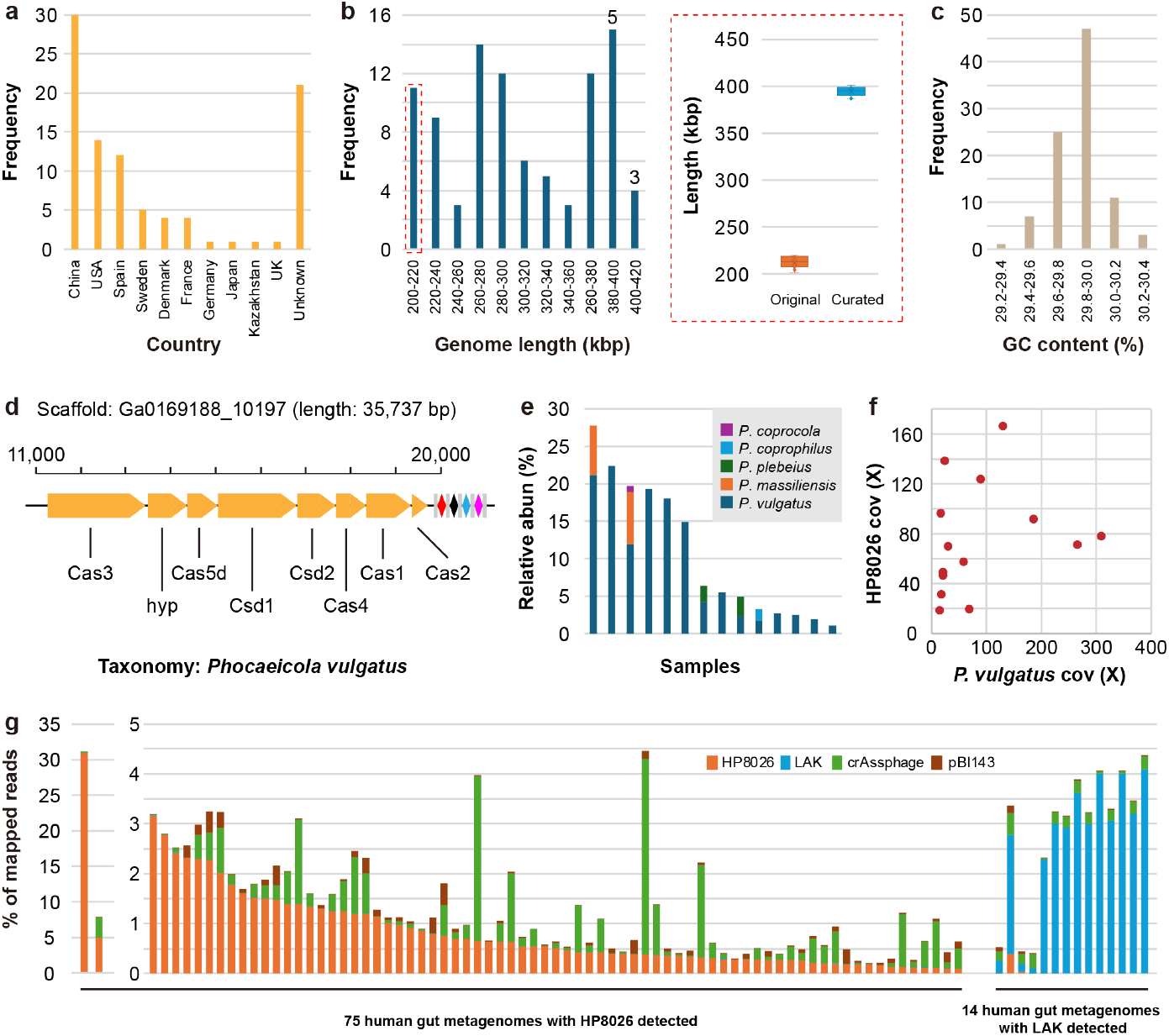
The general features and distribution of HP8026 clade. (a) The countries from where the HP8026 clade genomes were reconstructed. (b) The length distribution of HP8026 genomes. The number above the bar shows the count of complete genomes. The insertion shows the original length of genomes < 220 kbp and their curated length. (c) The GC contents of the HP8026 genomes. (d) CRISPR-Cas spacer matching suggested that *Phocaeicola vulgatus* is the host of HP8026 (see details in Supplementary Figure 5). (e) The relative abundance of *Phocaeicola* spp. and (f) the sequencing coverage of HP8026 and *P. vulgatus*, in 14 human gut metagenomic samples. (g) The relative abundance of mapped reads to HP8026, LAK, crAssphage, and pBI143 genomes in 75 human gut metagenomes with HP8026 genomes reconstructed and 14 human guts metagenomes with LAK phages detected.

Human guts are known to contain a high diversity of phages, such as crAssphage infecting *Bacteroides* spp. ^28^ and LAK phages infecting *Prevotella* spp. ^8,11^, and plasmids like pBI143 ^29^. HP8026 showed no significant sequence similarity to any of them. CRISPR spacer matching suggested that HP8026 likely infects *Phocaeicola vulgatus* (Figure 4d, Supplementary Figure 5), which was supported by co-occurrence analysis (Figures 4e and f). *Phocaeicola vulgatus* is a relative of *Bacteroides* (both belonging to Bacteroidaceae) and is usually abundant in human guts.

To reveal the distribution of HP8026 in human guts, we mapped metagenomic reads to the genomes of HP8026, crAssphage (comprising 168 groups at 80% similarity), LAK genomes, and pBI143. For the samples with HP8026 genomes reconstructed, HP8026 accounted for an average of 1.2% of the reads (up to 31.0%). In comparison, crAssphage and pBI143 had lower fractions (0.4% and 0.1%, respectively), and almost no LAK phages (Figure 4d). We also evaluated 14 human gut metagenomes with the LAK genome reconstructed and found HP8026 could be detected in only 2 samples. Using Pebblescout search ^30^, HP8026 was detected in over 2800 metagenomes (Supplementary Table 4). *De novo* assembly indicated that HP8026 may only be present in the human gut and related environments such as wastewater, with genomic divergences yet to be revealed (Supplementary Figure 6)

### The huge phage protein structure database, HPDB

We determined the best genetic code for the 7,295 non-redundant genomes, with 7,108 using code 11, and only 155 and 32 using code 15 and code 4, respectively. The protein-coding genes predicted from the non-redundant genomes were conducted for protein structure prediction and analyses. To have a reasonable representative of the genes and to reduce the number of genes for prediction, we performed a two-step clustering on the protein-coding genes before structure prediction. The 2,530,042 protein-coding genes were first clustered using mmSeqs2 to obtain 598,720 clusters (Supplementary Table 5). Then, the 36,971 mmSeqs2 clusters detected in ≥10 genomes were further clustered within each cluster using CD-HIT (see Methods). The representatives of CD-HIT clusters with ≥5 members were conducted for structure prediction (64,066 in total) using ColabFold ^31^. Approximately 80% (51,194) of the predicted structures had an average pLDDT (predicted local distance difference test) score of ≥70 (Figures 5a and b), representing 589,960 predicted proteins. These structures were clustered into 15,594 structure clusters using FoldSeek ^32^, with 64.7% of them (10,083) being singleton clusters.

**Figure 5.**
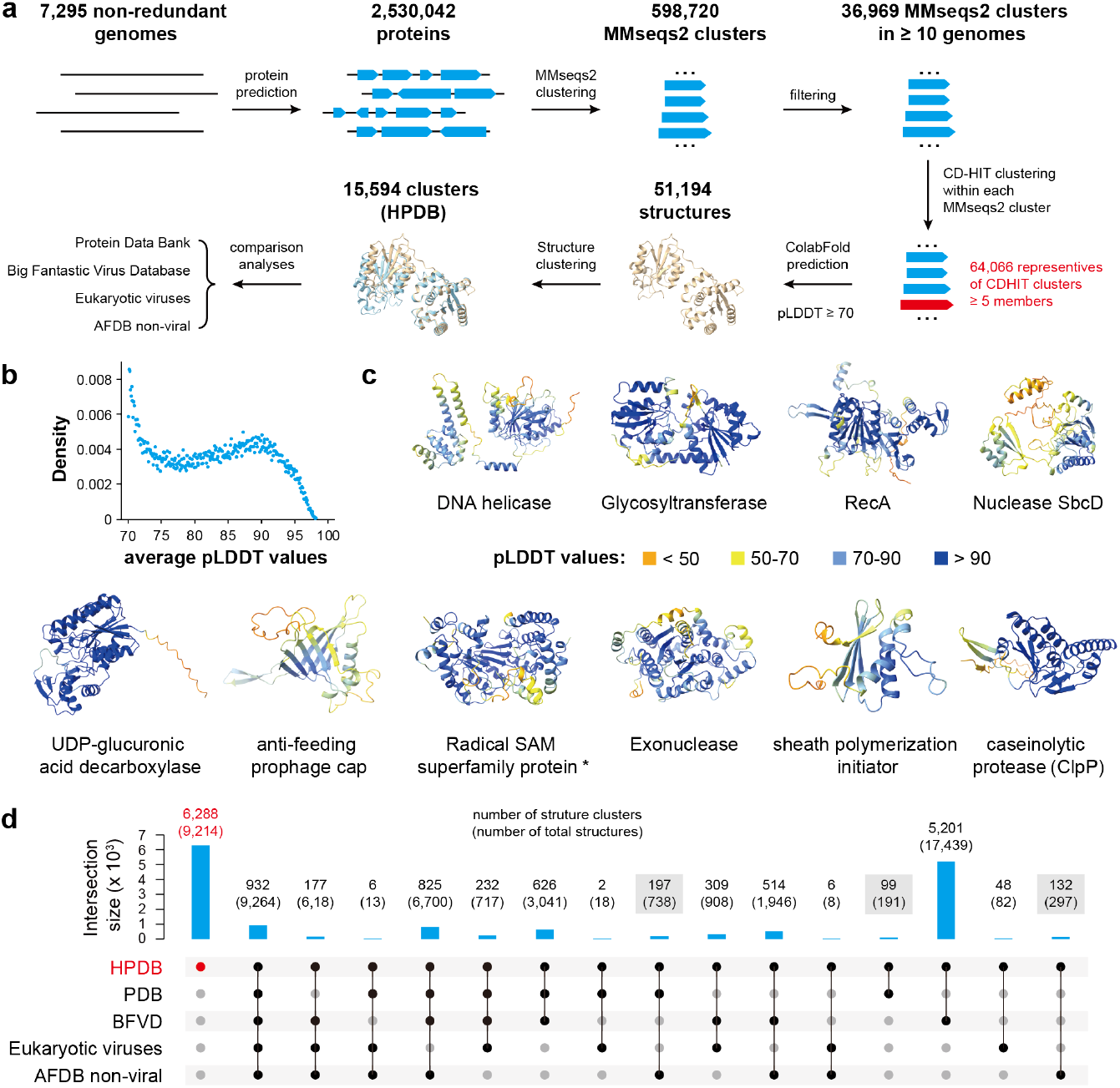
Protein structure prediction. (a) The pipelines of protein structure prediction. (b) The distribution of pLDDT values of the predicted structures. (c) The representatives of the top 10 most abundant structure clusters. The structure annotations were retrieved from the best PDB hits (with the lowest e-value), except the one marked with * that without PDB annotations. (d) The shared and unique HPDB protein structures. The HPDB structures were searched against the PDB, BFVD, Eukaryotic viruses structure database, and the non-viral AFDB using FoldSeek. The numbers of clusters present in huge phages while absent in other viral forms are highlighted with a gray background.

We investigated the nucleus-forming protein (i.e., chimallin) in huge phages to evaluate our protein structure prediction and clustering pipelines. We identified only one chimallin structure cluster (represented by huge_phage_762_291), which included 5 mmSeqs2 protein clusters containing 650 chimallin proteins (Supplementary Table 6). For 631 of them, their adjacent genes were for the DNA-directed RNA polymerase beta subunit, a conserved protein of nucleus-forming huge phages ^33^ (Supplementary Table 7), which was expected, suggesting that our analyses could efficiently link distantly related homologs.

As expected, all the top 10 most abundant structural clusters were related to basic functions (Figure 5c), including DNA processing (e.g., DNA helicase, RecA-like DNA repair protein, SbcD nuclease, and exonuclease), host manipulation mechanisms (e.g., glycosyltransferase and UDP-glucuronic acid decarboxylase), virion assembly (sheath polymerization initiator).

We compared the HPDB structures against those of PDB ^34^, BFVD (Big Fantastic Virus Database) ^35^, Eukaryotic viruses ^36^, and non-viral AFDB ^36^ (Figure 5d), and found that only ~6% were shared by all databases. Notably, ~40% of the HPDB structures were unique to huge phages, including those for basic functions such as terL, lysozyme, RNA polymerase, and others (Supplementary Table 8). Besides, the majority of the HPDB-specific structures were without any domain detected. These results indicated the structural uniqueness of huge phages.

### Protein folds present in huge phages while absent in other viral forms

We investigated the 428 HPDB structures (Figure 5d) that are not present in BFVD, which includes the structures of viral sequence representatives of the UniRef30 clusters ^35^, and the Eukaryotic viruses database ^36^, for huge phage structures not encoded by other viruses.

We found 102 antirestriction ArdC genes encoded by 101 genomes, with one genome having two copies. These huge phages were identified from various habitats, including marine, oil, freshwater, gut, and others, with a genome size of 203-677 kbp and a GC content of 22.3-66.2% (Supplementary Table 9).

ArdC was reported in the R388 plasmid to protect its ssDNA from endonucleases of the type I and III restriction and modification system during conjugation ^37–39^. ArdC contains a non-specific ssDNA binding N-terminal domain and a C-terminal metalloprotease domain. We aligned the 102 ArdC proteins against that of R388 and found only one without the key residues (Figure 6a). Structure clustering using FoldSeek grouped the ArdC proteins into two clusters, cluster 1 (including the one from R388, PDB id = 6i89) and cluster 2 comprising of 66 and 36 members, respectively (Figure 6b). By comparing against the R388 ArdC structure, the cluster 1 structures showed much lower global angstroms (Figure 6c). Interestingly, the cluster 1 could not be separated from cluster 2 by either ArdC phylogeny (Figure 6d) nor huge phage phylogeny (Supplementary Figure 7), highlighting the robustness of structure prediction and clustering to distinguish homologs while impossible at the sequence level. Moreover, the structural divergence despite phylogenetic overlap implies potential functional convergence or adaptive evolution of ArdC across distinct huge phage lineages. In summary, the detection of ArdC-encoded huge phages with various genome sizes in multiple habitats likely suggests that it is an important and efficient strategy for them to resist the antiviral mechanisms of their bacterial hosts.

**Figure 6.**
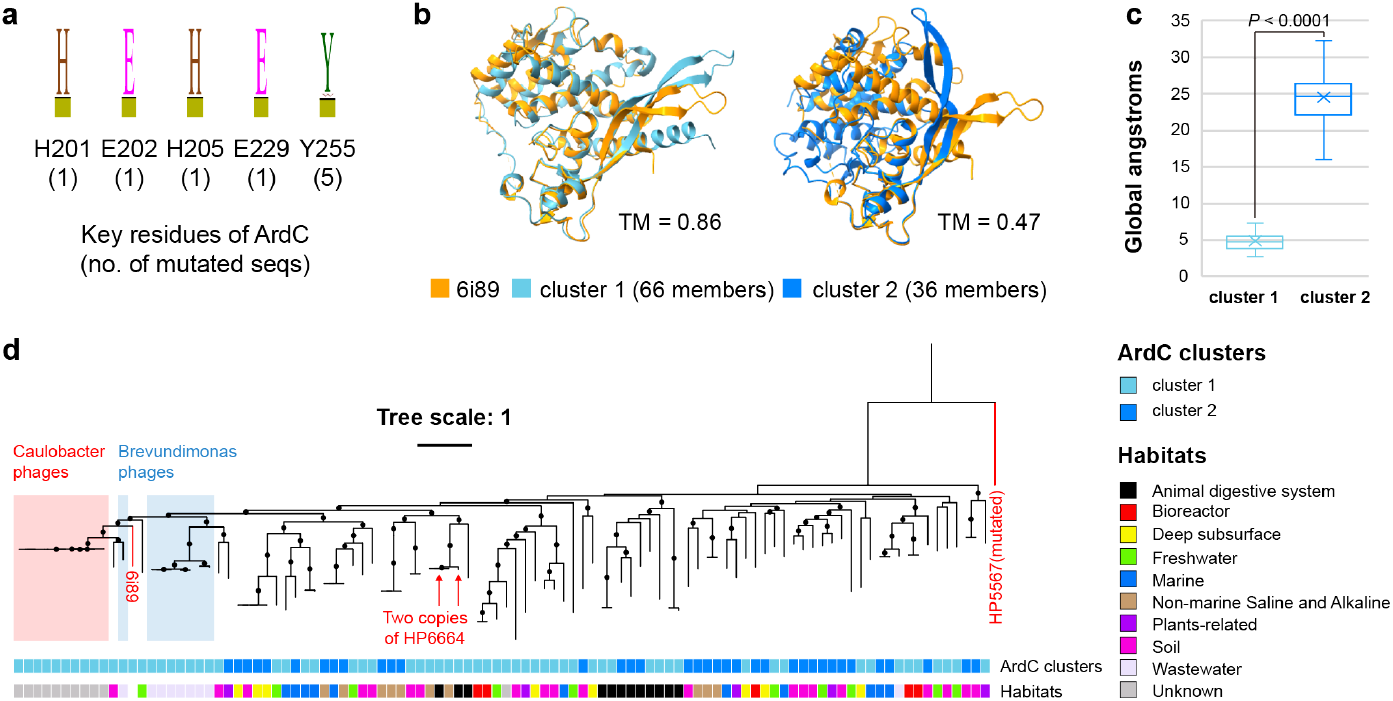
Huge phages encoded ArdC that may help them avoid DNA degradation. (a). The key residues of R388 plasmid ArdC were conserved in the majority of the ArdC proteins encoded by huge phages. All five key residues were mutated in the ArdC of HP5567. (b) The structure alignment of type 1 and 2 representatives against R388 plasmid ArdC (PDB id = 6i89). The TM-scores of the alignment are shown. (c) The global angstroms of all type 1 and 2 ArdC members to R388 plasmid ArdC. The statistical significance test was conducted using a two-tailed unpaired t-test. (d) The phylogeny of ArdC proteins. HP5567 ArdC was included as the outgroup. The two copies of ArdC from huge_phage_6664 (HP6664) were indicated by red arrows. The known hosts of hupe phages (Caulobacter and Brevundimonas) are shown with colored backgrounds. A black circle indicated a bootstrap value of ≥ 70.

Other interesting structures present in huge phages include the isocitrate lyase (EC 4.1.3.1) and malate synthase (EC 2.3.3.9), which are key to the glyoxylate cycle, a special variant of the tricarboxylic cycle (TCA) that allows utilization of two carbons compounds while glucose is absent or limited ^40^. The 98 isocitrate lyase and 22 malate synthase genes were identified in 98 huge phages, these two genes were adjacent to each other in 20 genomes, and one gene away in the other two genomes, suggesting their functional relatedness. Interestingly, though these 98 huge phages were from multiple habitats including freshwater, bioreactors, deep subsurface, marine, soil, and plant-related, the malate synthase encoded phages were only identified in the latter three habitats (Supplementary Table 10).

We also identified 248 genes for glutamine synthetase (GS; EC 6.3.1.2) in 247 huge phages, with 215 of them from freshwater and marine habitats (Supplementary Table 11). We speculate that the phage-encoded GS could enable them to directly modulate host nitrogen metabolism by synthesizing glutamine—a critical nitrogen carrier and precursor for nucleotides and amino acids, thus enhance their fitness under nitrogen-limited conditions ^41^. Additionally, phage-derived GS could destabilize host metabolic networks, redirecting nitrogen flux toward phage biomolecule synthesis. Such a trait would expand the paradigm of AMGs, highlighting their role in fine-tuning biogeochemical cycles (e.g., marine nitrogen turnover) and influencing host-phage coevolution.

Together, the abovementioned and many others that are not discussed in detail (Supplementary Table 6), suggested that huge phages may harbor many both basic (such as ArdC for avoiding DNA degradation) and metabolism-related functions, and thus are unique to other viral forms.

## Discussion

Due to the antibiotic resistance problem, phages that specifically infect bacteria, including animal and plant pathogens have been a hot topic in the field of microbial research ^42^. Huge phages are unique as they have large genome sizes and encode complex biological mechanisms that are usually absent in smaller phages ^3,16^, for example, the tiny type V CRISPR-Cas systems ^15^, the nucleus-like compartment that could help them escape from the immunity attack of their bacterial hosts ^18,20^. The past five years witnessed the enthusiasm towards huge phages, especially with the development of genome-resolved metagenomic and related analysis tools. Huge phages have been documented to be widespread in various habitats ^3^, more recent studies have revealed their diversity and evolution in undersampled environments such as glacier foreland soil ^43^. Notably, a more recent discovery of the first phages exceeding 800 kbp in genome size extended our knowledge of how big they could be ^6^.

To overcome the difficulties in the identification of huge phage sequences from the metagenomic assembly, which is also a challenge in identifying other viral sequences, we developed vRep, which combined the strength of available viral analysis tools such as geNomad ^26^, VirSorter2 ^23^, and CheckV ^25^. We emphasize the importance of obtaining viral sequences from the metagenomic assembly as reliably as possible, otherwise, all subsequent analyses will be impacted heavily including diversity and metabolisms ^44,45^. Using multiple viral identification tools and only counting the sequences predicted by more than one tool is the key, especially when working on huge datasets. One may argue that this may miss some novel viral sequences, which we agree with, and we suggest careful manual curation and inspection should be applied in such cases, such as Borgs ^46^. It should be mentioned that other viral identification tools could be used instead of any integrated into vRep, as each has advantages and disadvantages ^47^.

To overcome the challenge of lacking a comprehensive and curated huge phage reference dataset, we constructed HPGC. Compared to the huge number of published small phage genomes, up to tens to hundreds of thousands from a single study ^48,49^, we currently have a very limited source of huge phage genomes. We suggest that HPGC could be a robust resource for potential huge phage identification from the metagenomic assembly, especially when combined with single molecular sequencing ^50^, which could partially resolve the fragmentation issue in assembly with short reads.

HPGC accelerated the discovery of HP8026, which was detected in thousands of metagenomes via Pebblescout search and *de novo* assembly (Supplementary Figure 7), yet suggesting genomic divergence. Notably, we found some of the essential genes of HP8026, like large terminase, are highly similar among the HP8026 members, highlighting the potential for accurate identification in environmental samples, which could be the base of human fecal contamination strategy development. Besides, given the prevalence of HP8026 in human guts (Figure 4, Supplementary Table 4), further study should be focused on genomic diversity expansion to reveal their metabolisms and variations, as well as their potential relationship with human health. Furthermore, we encourage the identification of other overlooked but prevalent huge phage clades (Figure 3d), especially those detected in multiple environments that may offer an opportunity to study their adaptation strategies.

Another issue in huge phage analyses is the huge number of unannotated protein-coding genes, which hindered a better understanding of their specific biological mechanisms and their ecological significance ^6^. The establishment of HPDB started to addresses this challenge by providing structural insights into these enigmatic proteins (Supplemenatry Table 6). For instance, the identification of the plasmid-originated ArdC homologs in diverse huge phages highlights how structural clustering can uncover different functional analogs while impossible using sequence-based phylogeny (Figure 6d). Similarly, the discovery of glyoxylate cycle related enzymes and glutamine synthetase in huge phages from nutrient-limited habitats underscores the utility of integrating structural and metabolic annotations to reveal previously unidentified AMGs (Supplementary Tables 10 and 11). The structural uniqueness of ~40% of HPDB entries, particularly those lacking detectable domains, suggests that huge phages may encode entirely new functional modules yet to be cataloged (Figure 5d). Future studies combining protein structural predictions, heterologous expression, and targeted mutagenesis could validate these hypotheses. By bridging the annotation gap, HPDB not only illuminates the “dark matter” of phage proteomes but also positions huge phages as key players in microbial ecology and evolution, with implications for phage therapy and environmental engineering.

### The limitations of this study

Three principal limitations of this study should be acknowledged. First, although we compiled 7,295 non-redundant huge phage genomes, this collection may not fully encompass their rapidly expanding diversity. The accelerating pace of phage discovery through contemporary metagenomic studies suggests that our dataset, while substantial, likely represents a conservative estimate of existing genomic diversity. Second, all functional annotations were derived solely from computational predictions. The absence of experimental validation for any predicted protein function introduces inherent uncertainties. Third, structural predictions were performed only for a subset of protein-coding genes due to current computational limitations. While we have established a framework for systematic structure prediction using state-of-the-art tools to generate HPDB, the majority of genes await prediction and will be updated accordingly.

## Methods

### Development of vRep

The basic idea of vRep is to use multiple viral identification tools and their combined output information to identify viral genomes with high confidence. The input sequences were first evaluated by geNomad (parameter: end-to-end) ^26^ and VirSorter2 (parameters: --keep-original-seq --include-groups dsDNAphage,NCLDV,RNA,ssDNA,lavidaviridae) ^23^. The geNomad results were parsed using the “Conservative standard”, i.e., virus_score ≥ 0.80, n_hallmarks ≥ 1, mark_enrichment ≥ 1.5, and fdr ≤ 0.05. The VirSorter2 results were parsed to retain those with a score ≥ 0.5. If both geNomad and Virsorter2 identified a given sequence as virus-containing (provirus, or full-length virus) sequences, it would be retained for further analyses; otherwise, it would be flagged as “excluded (non-virus)”.

Then, CheckV ^25^ (parameter: end_to_end) was further utilized to analyze the virus-containing sequences to remove the host regions. Subsequently, the output files of “proviruses.fna” and “viruses.fna” from CheckV were combined (termed as “checkv.fa”). The CheckV provirus sequences will be conducted for another run of CheckV analysis to obtain the relevant information of the virus fragments.

The “checkv.fa” sequences (pure viral fragments) are evaluated by another run of VirSorter2 analyses with the parameters as follows, “--seqname-suffix-off --viral-gene-enrich-off --provirus-off --prep-for-dramv --include-groups dsDNAphage,NCLDV,RNA,ssDNA,lavidaviridae”. Subsequently, the genomes were filtered based on information from the CheckV and the second VirSorter2 analyses to retain those with (1) viral_gene >0 (i.e., Keep1), or (2) viral_gene =0 and (host_gene = 0 or score >=0.95 or hallmark >2) (i.e., Keep2), following a widely used online procedure ^51^.

To check the replication issue that may arise in the de novo assembly, the retained sequences will be checked with a BLASTn search, and for a given sequence, if its first half and second half share 50% of their length with ≥ 99% nucleotide sequence similarity will be flagged.

In the final step, a summary will be generated, including information from CheckV, VirSorter2, and geNomad and duplication issues for each virus-containing sequence. The users could then screen their sequences for further analysis.

### Collection and evaluation of potential huge phage genomes using vRep

Published or public potential huge phage genomes were collected from IMG/VR and NCBI databases and publications via the downloading links or accession numbers if provided.

#### IMG/VR

The IMG/VR database was filtered to retain the viral genomes (1) with a minimum length of 200 kbp, (2) not assigned as “Megaviricetes” (via the corresponding IMG/VR mapping file), and (3) with the corresponding sample is not under the JGI Data Utilization Status of “Restricted”, which obtained a total of 5,181 sequences.

#### NCBI

The NCBI phage genomes with a minimum length of 200 kbp were collected with the keyword “phage”, and the “Sequence length” was set from “200000” to “1000000”, which retrieved a total of 954 genomes on September 3rd, 2024 (“NCBI_phage_200kbp”). After evaluation by vRep, 817 were identified as huge phage genomes. To avoid missing any genomes of huge phages somehow named “virus” in NCBI, we also searched “virus” with the same length range, and retrieved a total of 943 genomes (“NCBI_virus_200kbp”). Among them, 74 were determined as huge phage genomes using vRep, and 68 were already included in the “NCBI_phage_200kbp” set. Thus, a total of 6 more huge phage genomes were obtained. In summary, 823 huge phage genomes were obtained from NCBI (“NCBI_huge_phages”). Local genome comparison indicated that all the previously reported isolated huge phages ^5^ were all included in “NCBI_huge_phages”.

#### Publications

The databases published before August 31, 2024, were downloaded according to the descriptions in the corresponding publications (if available). A detailed description of the publications and databases can be found in Supplementary Table 2. The viral genomes were filtered to retain those with a minimum length of 200 kbp, followed by vRep evaluation to screen for huge phage genomes. We also retrieved some metagenomic datasets sequenced with the 3rd generation sequencing technology such as Nanopore. For example, for the Nanopore sequences (passed default Guppy quality control) reported recently ^52^ were provided by the authors, those with a minimum length of 200 kbp were evaluated using vRep, and 203 of them were identified as huge phage genomes.

### Evaluation of huge phage genomes from publications

The genomes from publications focusing on phages were evaluated to identify the common misidentification issues in huge phage analyses ^3,9,10,14,17,53–57^. In detail, the phage genomes with a minimum length of 200 kbp were downloaded and evaluated by vRep for potential misidentification issues. From each study, the number of sequences that were assigned to the misclassification cases was summarized (Supplementary Table 1).

### Alternative coding analyses

Prodigal (single mode) was used to predict genes on each genome using genetic codes 4, 11, and 15, as previously described ^58^. The coding density of each genome was calculated by adding the total number of coding bases and dividing by the total genome length. The genomes with a coding density increase of ≥5% in code 4 or code 15 relative to code 11 were assigned that genetic code.

### Genome clustering

To remove redundant genome sequences that may be reported from different studies or different assemblies, all the identified huge phage genomes were clustered using the rapid genome clustering approach provided by CheckV (available at https://bitbucket.org/berkeleylab/checkv/src/master/) ^25^, with the parameters set as “--min_ani 100 --min_tcov 100”, to obtain the non-redundant genome set. These non-redundant genomes were further clustered into species-level groups using the same approach, with the parameters set as “--min_ani 95 --min_tcov 85”.

### Identification and analyses of HP8026

We attempted to determine how many species-level groups could be further clustered using lower sequence similarity, i.e., 80%, and found that HP8026 was among the few of them.

To curate the low-quality HP8026 clade genomes, the corresponding paired-end read files were downloaded from NCBI and assembled using metaSPAdes ^59^. The assembled scaffolds were then searched against the HP8026 genomes using BLASTn for HP8026-related scaffolds. The target scaffolds were manually curated via read mapping, scaffold extension, and reassembly, as previously described ^60^.

To compare the HP8206 genomes against crAssphage (145-192 kbp) and LAK phages (408-660 kbp), their genomes were downloaded according to the corresponding publications ^8,11,28^ and searched using BLASTn using an e-value threshold of 1e^-100^.

To evaluate the distribution of HP8026 huge phages, the paired-end reads from the corresponding samples from where the huge phage genomes were constructed were downloaded from NCBI, and mapped to the HP8026, crAssphage, LAK phage, and pBI143 genomes using Bowtie2 ^61^ with default parameters. The Shriksam tool (https://github.com/bcthomas/shrinksam) was used to exclude the unmapped pairs from the sam file, which was subsequently sorted and transferred to the bam format file using Samtools ^62^. The number of mapped reads to each genome across samples was generated by CoverM version 0.7.0 (“contig” mode) (https://github.com/wwood/CoverM) using the “count” method with the parameters set as “--min-read-aligned-percent 95 --min-read-percent-identity 90 --contig-end-exclusion 75 -m count”.

To evaluate the protein source and metabolic potentials of HP8026, the predicted protein-coding genes were first clustered using CD-HIT version 4.8.1 ^63^ with parameters of “-c 0.9 -aL 0.8 -aS 0.8”. The cluster representatives were conducted for protein structure predictions using ColabFold (downloaded on 22 Nov 2024) ^31,64^, with the parameters as follows, “--num-recycle 3, --use-gpu-relax, --amber, --stop-at-score 70”. For structure annotation, the predicted structures with a pLDDT score of ≥ 70 were searched against the protein data bank (PDB) ^34^ using FoldSeek 9.427df8a ^32^ with the “easy-search” command (-c 0.5 --cov-mode 0). Only those hits with a TM score ≥ 0.5 (“alntmscore”) were retained for further analyses. As a result, 38,574 predicted proteins were clustered into 1,831 CD-HIT clusters, whose representatives were conducted for structure predictions, and 189 of them were with PDB annotations (Supplementary Table 5).

### Protein family clustering and protein structure prediction

The protein sequences predicted from all the 7,295 non-redundant huge phage genomes were first clustered using MMseqs2 version 15.6f452 ^65^. Specifically, an all-vs.-all search was performed using “-s 7.5 -e 0.001 --min-seq-id 0.2 -c 0.7 --cov-mode 0 --max-seqs 50000 --cluster-mode 0”.

To reduce the number of sequences for protein structure prediction, we further picked up the MMseqs2 clusters presented in ≥ 10 non-redundant genomes. Then, a second clustering was performed within each of the picked MMseqs2 clusters using CD-HIT version 4.8.1 ^63^, with the parameters “-c 0.7 -aL 0.5 -aS 0.5 -G 0”. The representatives of CD-HIT clusters with ≥ 5 members were extracted for structure prediction using ColabFold (downloaded on 22 Nov 2024) ^31,64^. The structure prediction parameters were as follows, “--num-recycle 3, --use-gpu-relax, --amber, --stop-at-score 70”.

### Protein structure clustering and comparison

The rank 001 models predicted structures with an average ColabFold pLDDT score of ≥ 70 were clustered using FoldSeek 9.427df8a ^32^ “easy-cluster” with the parameters of “-c 0.7 --cov-mode 0 --tmscore-threshold 0.5”. The representative structure of each structure cluster was then searched against (1) the protein data bank (PDB) ^34^, (2) the Big Fantastic Virus Database (BFVD) ^35^, (3) the eukaryotic viruses structure database^36^, and (4) the initial release of AlphaFold database that contains more than 500,000 proteins from eukaryotes, bacteria, and archaea without virus (AFDB non-viral) ^66^, using FoldSeek 9.427df8a ^32^ with the “easy-search” command (-c 0.7 --cov-mode 0). Only those with a TM score ≥ 0.5 (“alntmscore”) were thought to be reliable matches.

### Identification of large terminase proteins

The predicted proteins were searched against a collection of huge phage large terminase proteins from NCBI and a previous publication ^3^ using BLASTp ^67^ (e-value threshold, 1e-100). All the proteins in a given MMseqs2 cluster with one or more hits were extracted for protein structure prediction using ColabFold, as described above. A total of 7,009 potential large terminase proteins were predicted with a minimum pLDDT value of ≥70. The predicted structures were clustered using FoldSeek ^32^, requiring the alignment to consist of at least 70% query and target coverage (i.e., -c 0.7 --cov-mode 0).

### Protein structure analyses

The protein structures of interest were manually inspected using ChimeraX version 1.8 ^68^. The alignment and comparison of R388 ArdC (6i89) against those identified in huge phages were performed using the Needleman-Wunsch alignment algorithm and the BLOSUM-62 similarity matrix, with the gap open (HH/SS/other) as 18/18/6 and gap extend as 1. The visualization of the structure alignment was also conducted using ChimeraX version 1.8.

### Phylogenetic analyses

For the large terminase and ArdC protein sequences, the alignment was performed using MUSCLE v3.8.31 ^69^ and filtered using trimAl v1.4.rev22 ^70^ to remove those columns comprising ≥90% gaps. The phylogenetic trees were constructed using IQ-TREE ^71^ with the parameter “-bb 1000, -m LG+G4”.

