## Supplementary Figures for "The hidden prevalence and unique protein folds of huge phages"

for

LinXing Chen (陈林兴) [1,*]

[1] State Key Laboratory of Advanced Environmental Technology, the Department of Environmental Science and Engineering, University of Science and Technology of China, Hefei, China, 230026

[*]Corresponding author:

Linxing Chen,

**Acknowledgements**

We thank Dr. Zhengshuang Hua for providing computational resources, and Dr. Xingxing Shen for helpful discussion in the initial stage of this work. We thank Lauren Lui for providing the quality-controlled Nanopore sequences. We thank Antonio Pedro Camargo for guiding the usage of IMG/VR v4 sequences regarding their open or restricted access. We thank the Super Computing Center at the University of Science and Technology of China for its support of ColabFold analyses. This work was financially supported by the grant funding of KY2400000036.

**Author contributions**

L.X.C. conceived this study, performed all the analyses, and drafted and finalized the manuscript.

**Competing interests**

The author declares no competing interests.

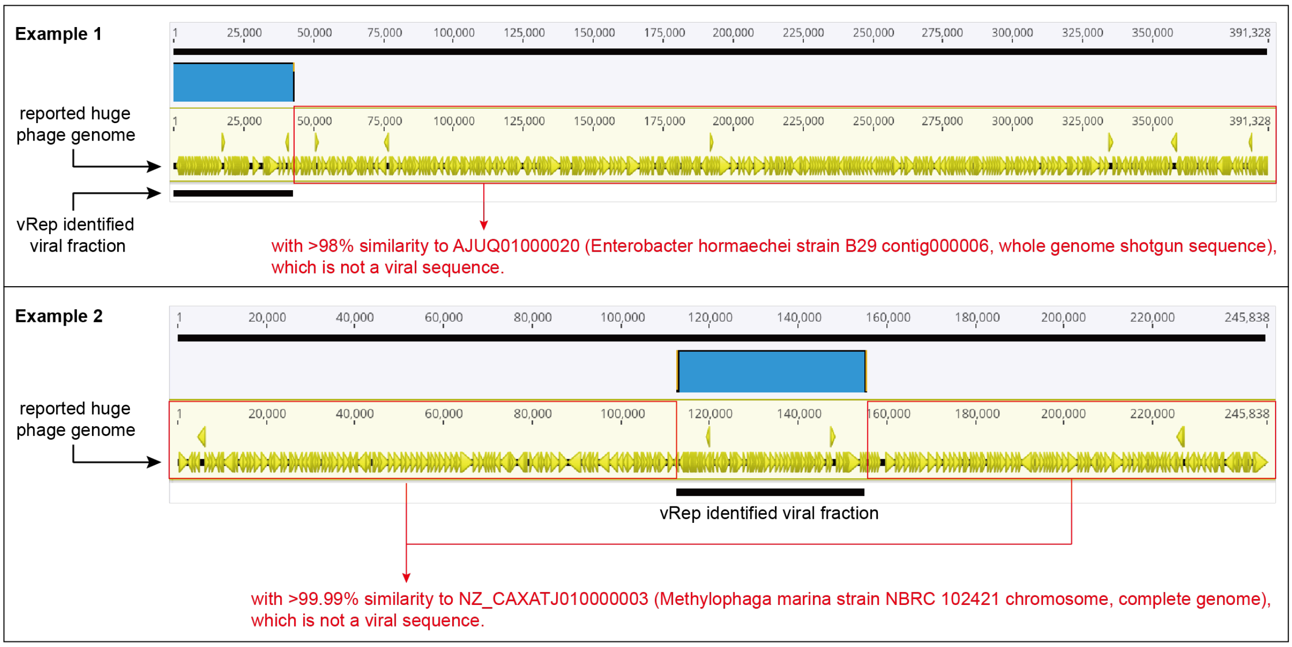


**Supplementary Figure 1 | Two more examples of huge phage misidentification case I issue.** In these examples, the metagenomic sequence with both phage and bacterial host genomic fractions were misidentified as pure phage sequences, thus assigned as huge phage genomes once the length exceeding 200 kbp. See **Figure 2c** in the main text for another example.

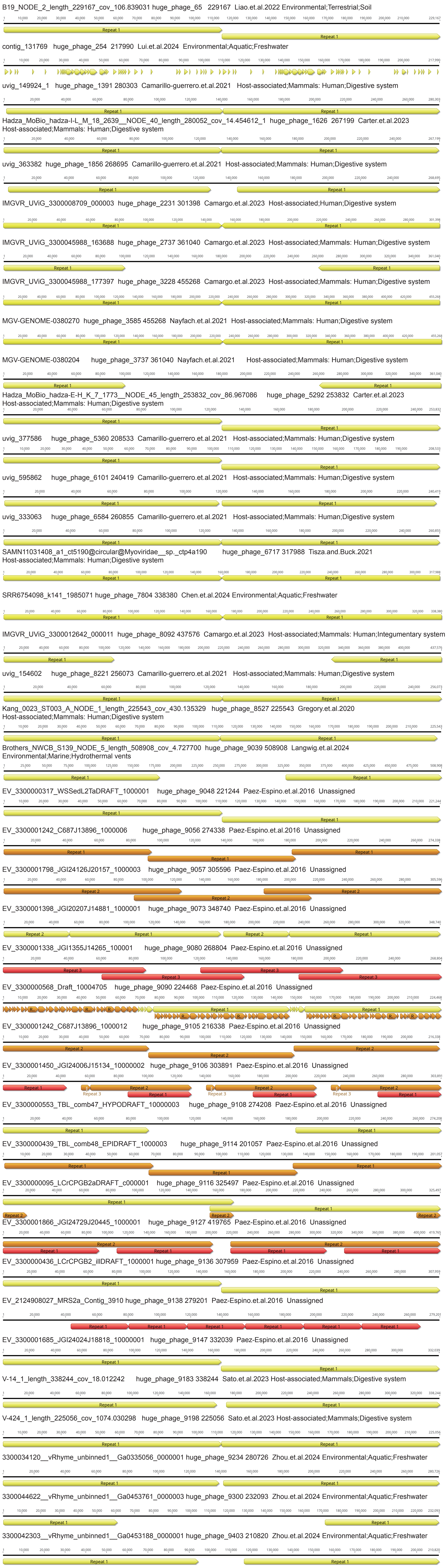
B19_NODE_2_length_229167_cov_106.839031 huge_phage_65 229167 Liao.et.al.2022 Environmental;Terrestrial;Soil
contig_131769 huge_phage_254 217990 Lui.et.al.2024 Environmental;Aquatic;Freshwater
uvig_149924_1 huge_phage_1391 280303 Camarillo-guerrero.et.al.2021 Host-associated;Mammals: Human;Digestive system
Hadza_MoBio_hadza-I-L_M_18_2639__NODE_40_length_280052_cov_14.454612_1 huge_phage_1626 267199 Carter.et.al.2023 Host-associated;Mammals: Human;Digestive system
uvig_363382 huge_phage_1856 268695 Camarillo-guerrero.et.al.2021 Host-associated;Mammals: Human;Digestive system
IMGVR_UViG_3300008709_000003 huge_phage_2231 301398 Camargo.et.al.2023 Host-associated;Human;Digestive system
IMGVR_UViG_3300045988_163688 huge_phage_2737 361040 Camargo.et.al.2023 Host-associated;Mammals: Human;Digestive system
IMGVR_UViG_3300045988_177397 huge_phage_3228 455268 Camargo.et.al.2023 Host-associated;Mammals: Human;Digestive system
MGV-GENOME-0380270 huge_phage_3585 455268 Nayfach.et.al.2021 Host-associated;Mammals: Human;Digestive system
MGV-GENOME-0380204 huge_phage_3737 361040 Nayfach.et.al.2021 Host-associated;Mammals: Human;Digestive system
Hadza_MoBio_hadza-E-H_K_7_1773__NODE_45_length_253832_cov_86.967056 huge_phage_5292 253832 Carter.et.al.2023 Host-associated;Mammals: Human;Digestive system
uvig_377586 huge_phage_5360 208533 Camarillo-guerrero.et.al.2021 Host-associated;Mammals: Human;Digestive system
uvig_595862 huge_phage_6101 240419 Camarillo-guerrero.et.al.2021 Host-associated;Mammals: Human;Digestive system
uvig_333063 huge_phage_6584 260855 Camarillo-guerrero.et.al.2021 Host-associated;Mammals: Human;Digestive system
SAMN11031408_a1_ct5190@circular@Myoviridae__sp._ctp4a190 huge_phage_6717 317988 Tisza.and.Buck.2021 Host-associated;Mammals: Human;Digestive system
SRR6754098_k141_1985071 huge_phage_7804 338380 Chen.et.al.2024 Environmental;Aquatic;Freshwater
IMGVR_UViG_3300012642_000011 huge_phage_8092 437576 Camargo.et.al.2023 Host-associated;Mammals: Human;Integumentary system
uvig_154602 huge_phage_8221 256073 Camarillo-guerrero.et.al.2021 Host-associated;Mammals: Human;Digestive system
Kang_0023_ST003_A_NODE_1_length_225543_cov_430.135329 huge_phage_8527 225543 Gregory.et.al.2020 Host-associated;Mammals: Human;Digestive system
Brothers_NWCB_S139_NODE_5_length_508908_cov_4.727700 huge_phage_9039 508908 Langwig.et.al.2024 Environmental;Marine;Hydrothermal vents
EV_3300000317_WSSedL2TaDRAFT_1000001 huge_phage_9048 221244 Paez-Espino.et.al.2016 Unassigned
EV_3300001242_C687J13896_1000006 huge_phage_9056 274338 Paez-Espino.et.al.2016 Unassigned
EV_3300001798_JGI24126J20157_1000003 huge_phage_9057 305596 Paez-Espino.et.al.2016 Unassigned
EV_3300001398_JGI20207J14881_1000001 huge_phage_9073 348740 Paez-Espino.et.al.2016 Unassigned
EV_3300001338_JGI1355J14265_100001 huge_phage_9080 268804 Paez-Espino.et.al.2016 Unassigned
EV_3300000568_Draft_10004705 huge_phage_9090 224468 Paez-Espino.et.al.2016 Unassigned
EV_3300001242_C687J13896_1000012 huge_phage_9105 216338 Paez-Espino.et.al.2016 Unassigned
EV_3300001450_JGI24006J15134_10000002 huge_phage_9106 303891 Paez-Espino.et.al.2016 Unassigned
EV_3300000553_TBL_comb47_HYPODRAFT_10000003 huge_phage_9108 274208 Paez-Espino.et.al.2016 Unassigned
EV_3300000439_TBL_comb48_EPIDRAFT_1000003 huge_phage_9114 201057 Paez-Espino.et.al.2016 Unassigned
EV_3300000095_LCrCPGB2aDRAFT_c000001 huge_phage_9116 325497 Paez-Espino.et.al.2016 Unassigned
EV_3300001866_JGI24729J20445_1000001 huge_phage_9127 419765 Paez-Espino.et.al.2016 Unassigned
EV_3300000436_LCrCPGB2_illDRAFT_1000001 huge_phage_9136 307959 Paez-Espino.et.al.2016 Unassigned
EV_2124908027_MRS2a_Contig_3910 huge_phage_9138 279201 Paez-Espino.et.al.2016 Unassigned
EV_3300001685_JGI24024J18818_10000001 huge_phage_9147 332039 Paez-Espino.et.al.2016 Unassigned
V-14_1_length_338244_cov_18.012242 huge_phage_9183 338244 Sato.et.al.2023 Host-associated;Mammals;Digestive system
V-424_1_length_225056_cov_1074.030298 huge_phage_9198 225056 Sato.et.al.2023 Host-associated;Mammals;Digestive system
3300034120__vRhyme_unbinned1__Ga0335056_0000001 huge_phage_9234 280726 Zhou.et.al.2024 Environmental;Aquatic;Freshwater
3300044622__vRhyme_unbinned1__Ga0453761_0000003 huge_phage_9300 232093 Zhou.et.al.2024 Environmental;Aquatic;Freshwater
3300042303__vRhyme_unbinned1__Ga0453188_0000001 huge_phage_9403 210820 Zhou.et.al.2024 Environmental;Aquatic;Freshwater

**Supplementary Figure 2 | The genome duplication issue (case IV) that raised during metagenomic assembly.** In these examples, a large fraction of a given genome was somehow present in two or more copies, thus could make a small genome much longer.

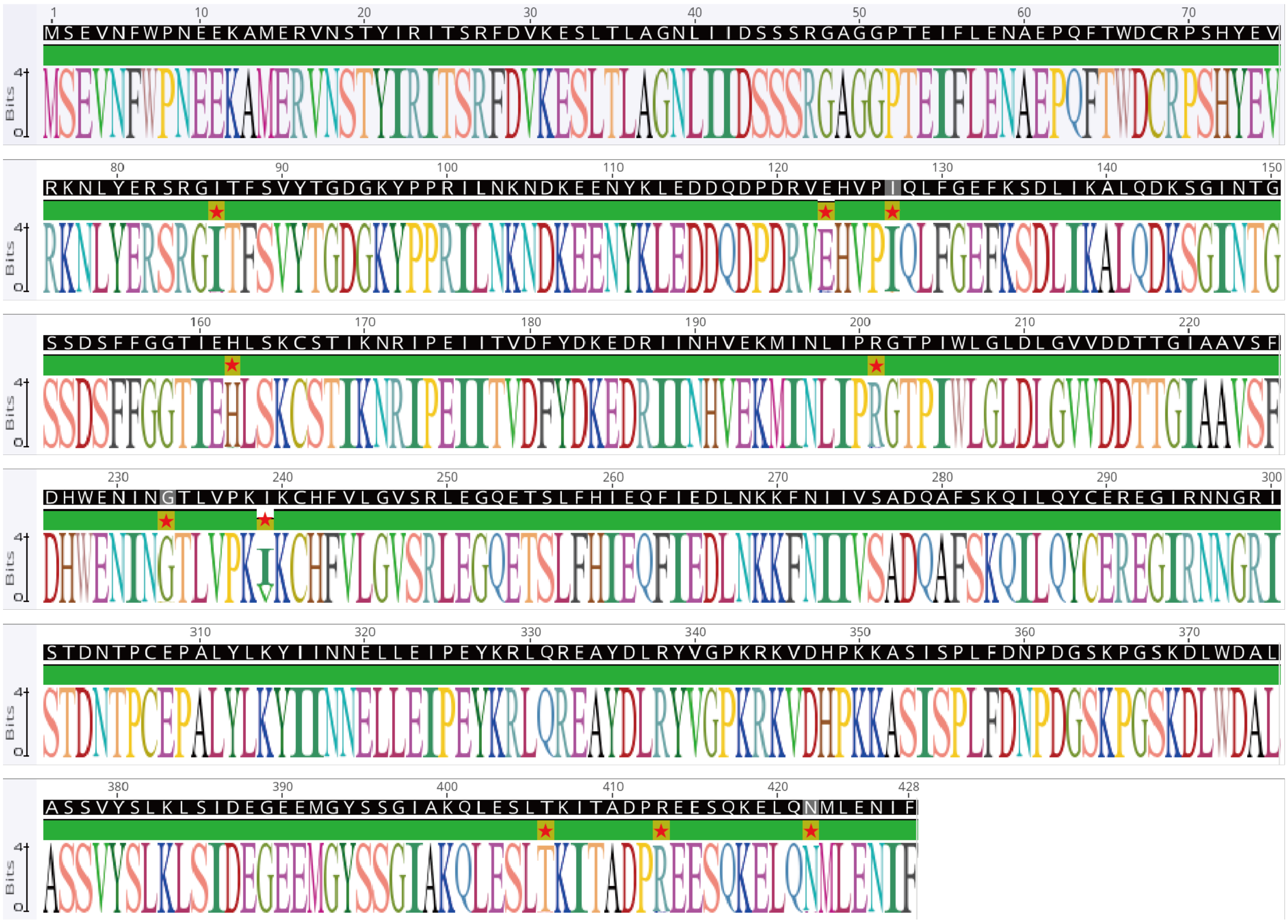

**Supplementary Figure 3 | Sequence logo shows the conservation of the large terminase proteins encoded by HP8026 clade genomes.** Due to the incompleteness of some of the 94 genomes, only 67 genomes were identified with the large terminase genes and thus included in the analyses. The red stars (10 in total) show the positions with divergences.

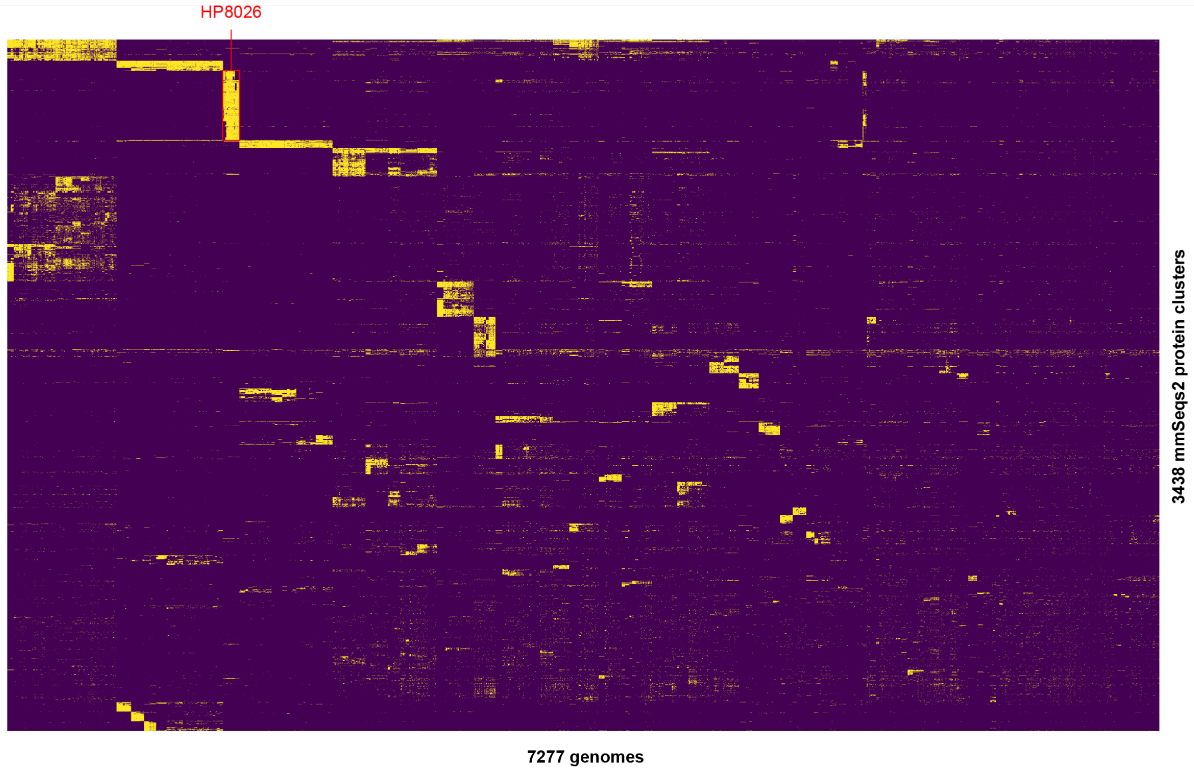


**Supplementary Figure 4 | The clustering of mmSeq2 protein clusters distributed in the non-redundant huge phage genomes.** The clustering was performed using the of method of 'ward' and the metric of 'euclidean'. Only the mmSeqs2 protein clusters present in at least 1% (i.e., 73 genomes) of all the 7,295 non-redundant huge phage genomes were included, which resulted in 7,277 genomes and 3,438 protein clusters. The HP8026 clade genomes were highlighted with a rex box.

Sample name: Human fecal microbial communities from newborn in Denmark - 128_B
NCBI SRA accession: ERR525705
Isolation: Human feces from Newborn
Country: Denmark
Scaffold ID: Ga0169188_10197
Scaffold length: 35,737 bp
CRISPR-Cas repeat sequence: GTCACACCCTGCGTGGGTGTGTGGATTGAAAC

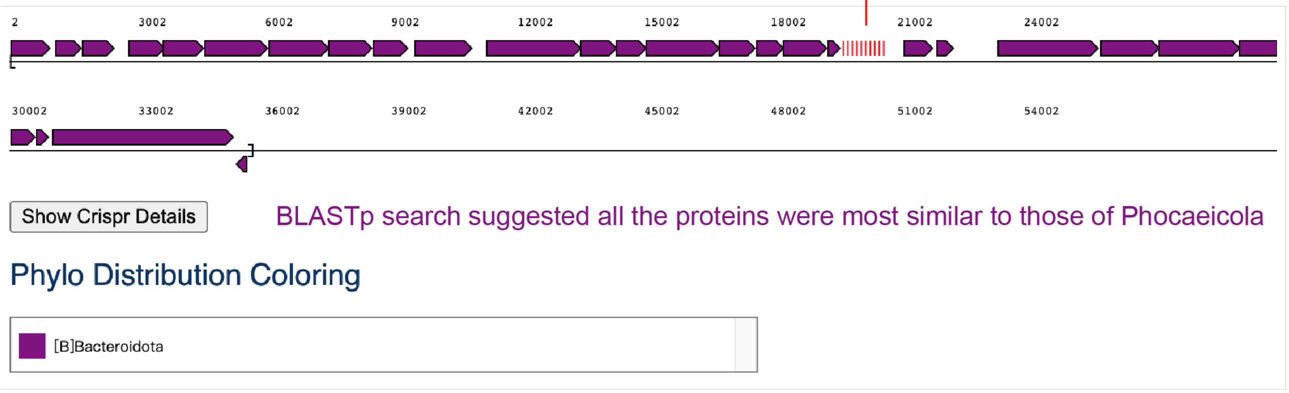


Spacer: AGTGATAGTAATATAGATAGTTTAGCGGTTGATAT
HP8026: AGTGATAGTAATATAGATAGTTTAGCGGTAGATAT → identity = 34/35 = 97.1%

→ part of a gene encoding a hypothetical protein

**Supplementary Figure 5 | CRISPR-Cas spacer targeting analysis showed that *Phocaeicola* is a potential host of HP8026 members**. iPHoP analysis was first performed on all HP8026 genomes, then the CRISPR-Cas spacer matching results were manually confirmed, including the detection of Cas proteins, the taxonomic assignment of the CRISPR-Cas system carrying scaffolds, to exclude potentially misleading conclusion.

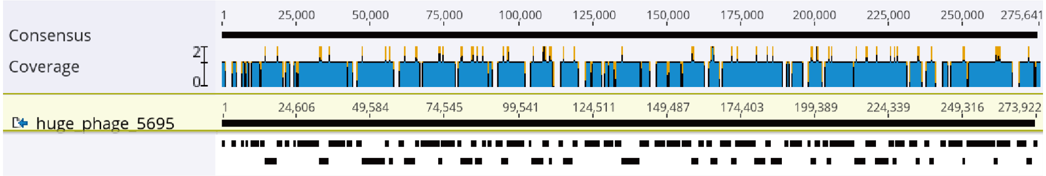


ERR4682349 Global surveillance of antimicrobial resistance - batch November 2017

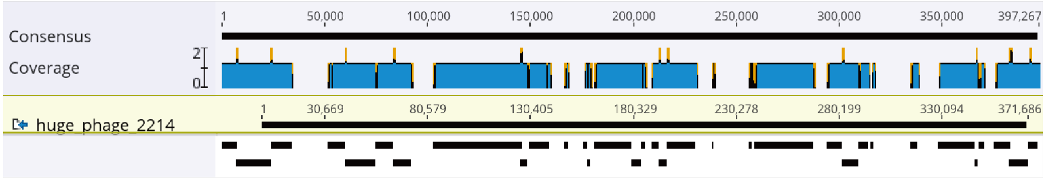


SRR10682561 viral metagenome Metagenome

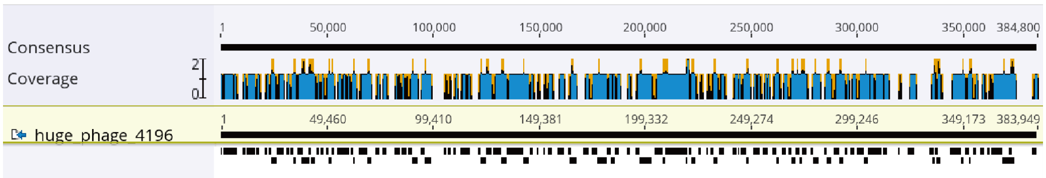


SRR8715493 Israel Pilot-Scale Hospital Wastewater Treatment System Raw sequence reads

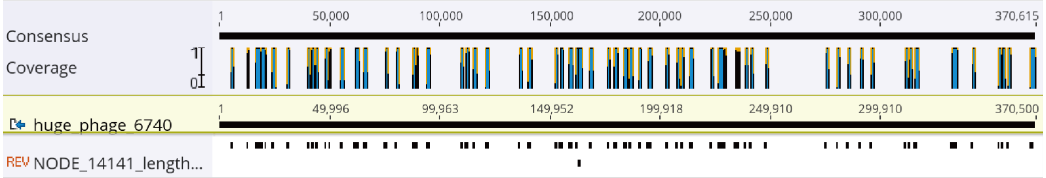


SRR15322609 Mouse intestinal phages-Health and alcohol feeding

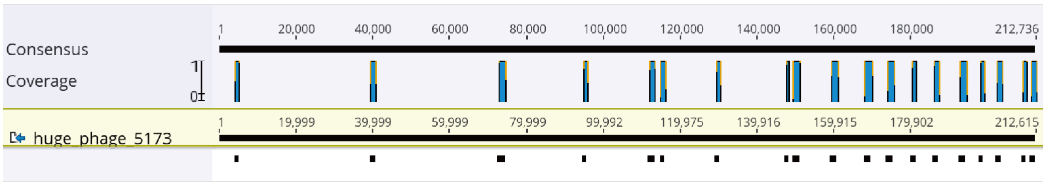


SRR13765886 gut microbiomes of dogs and wolves

**Supplementary Figure 6 | The alignment of Pebblescout search retrieved metagenomic assembly to HP8026 genomes.** The paired-end reads of five Pebblescout search retrieved samples with lower %coverage values (Supplementary Table 4) were de novo assembled, and the scaffolds with a minimum nucleotide similarity of 90% over a minimum length of 1000 bp (determined by BLASTn) to HP8026 genomes were aligned to the corresponding genomes and shown.

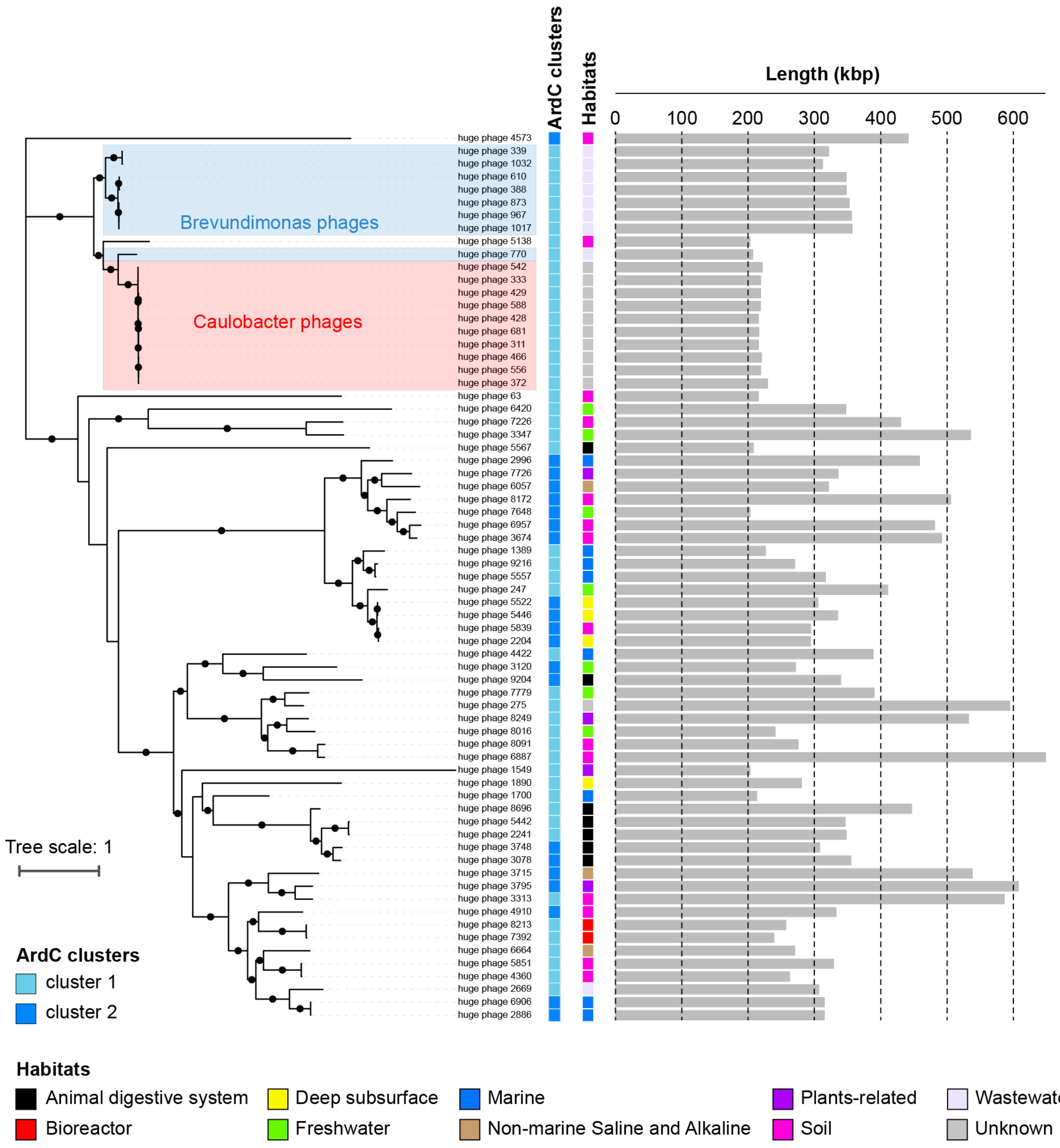


**Supplementary Figure 7 | The phylogeny of ArdC-encoding huge phages based on the large terminase (terL) protein sequences.** A total of 68 genomes were included. The name of the huge phage encodes a ArdC with the conserved key residues mutated was highlighted in red. The colored strips of the inner circle indicate the ArdC type, and the colored strips of the outer circle show the habitats. The gray bars show the length of the genomes. The huge phages with known hosts are highlighted with colored background. The black circle indicates a bootstrap value of ≥ 70.
